# Optimization of Tissue Clearing Methods and Imaging Conditions for 3D Visualization of the Vasculature of the Adult Murine Knee

**DOI:** 10.1101/2025.06.26.661802

**Authors:** Azeez O. Ishola, Athira Pillai, Taeyong Ahn, RE-JOIN Consortium, Chih-Wei Hsu, Brendan Lee, Nele Haelterman, Joshua D. Wythe

**Author notes:** Corresponding author To whom correspondence should be addressed: Joshua D. Wythe, Ph.D. University of Virginia School of Medicine Charlottesville, VA 22903.

## Abstract

The vasculature is an essential regulator of tissue oxygenation, metabolism, and immune surveillance under developmental and homeostatic conditions, as well as in regeneration and disease. Within the musculoskeletal system, blood vessels are involved in bone development and resorption, and they also mediate the inflammatory processes that contribute to diseases affecting the joints, including osteoarthritis. Historically, visualization of vascular networks in joints has been limited to conventional histological approaches due to technical limitations and the inherit structure of the skeletal muscle and bones. Recent advances in optical tissue clearing technologies and 3D fluorescence imaging approaches enabling three-dimensional analysis in whole, intact tissues offer an entrée to interrogate musculoskeletal development and disease at a cellular resolution. However, these techniques were not originally developed to image hard mineralized tissues, such as the femur and tibias. To circumvent these challenges, we have optimized tissue decalcification conditions and systematically compared aqueous- and solvent-based tissue clearing approaches to establish an optimal pipeline for clearing and imaging of mineralized tissues with superior resolution of the vasculature. Collectively, this work shows that optical clearing and lightsheet microscopy presents a viable method for generating high-resolution images of the murine hindlimb vasculature, with potential applications in aging and disease modeling.

## INTRODUCTION

The World Health Organization estimates that more than 1.71 billion people worldwide suffer from musculoskeletal disease^1,2^. From rheumatoid arthritis to gout, osteoporosis, spinal disorders, lower back pain and severe trauma, more than 150 unique musculoskeletal diseases impact people globally, leading to extensive health care expenditures^3^. Given the number of tissues that make up the musculoskeletal system – including the bones, muscles, ligaments, tendons, cartilage and more – the prevalence of these ailments is perhaps unsurprising. Already the leading cause of physical disability, as the world population ages, and risk factors increase, the number of individuals suffering from these diseases are only predicted to rise^4^. To combat the increased incidence and severity of these diseases requires a deeper understanding of the tissues comprising the musculoskeletal system and how they interact in both health and disease, particularly osteoarthritis (OA).

OA is a complex and chronic degenerative disease affecting the whole synovial joint, characterized by structural defects and damage to the articular cartilage, loss of subchondral bone, tissue hypertrophy, increasing vascularity of the synovium, synovitis and immune cell recruitment and activation, fibrosis, and instability of the ligaments and tendons^5,6^. A multifaceted joint disease, OA involves systemic modifications and numerous comorbidities that influence overall health, including cardiovascular disease, obesity, type 2 diabetes, and more^7^. With over 500 million cases worldwide, OA is estimated to be the fourth leading cause of disability, with an enormous socio-economic burden globally due health care costs, loss of work, and early retirement^8^. Currently, there is no cure for OA, and these patients account for 55.3% of all opioid prescriptions in the US, resulting in about $14 billion in lifetime opioid-related societal costs ^9^. Accordingly, a more comprehensive understanding of tissue dynamics during OA initiation and progression are needed to more effectively treat this devastating disease.

As a weight-bearing structure subjected to lifelong mechanical stressors and a frequent pathological changes, the knee accounts for more than 260 million cases of OA ^10–12^. Historically, studies of OA in the knee relied on thin serial sections followed by immunostaining and imaging. However, mechanical sectioning is labor intensive, often destructive, and provides only a single snapshot, in one plane, of what is happening within and around a joint or bone. Alternatively, more recent studies have employed 3D imaging modalities, such as ultrasound or micro-computed tomography (micro-CT) ^13^. However, increasing our understanding of OA pathogenesis necessitates defining anatomical maps of tissue relationships and cellular circuits in healthy and diseased states through simultaneous imaging of the bones and associated tissues of the knee joint in three-dimensional space at a cellular resolution, which is beyond what is capable with imaging methods such as micro-CT and MRI. Serendipitously, advances over the last 15 years or so in tissue clearing ^14,15^ and light microscopy ^16^ have the potential to revolutionize the study of musculoskeletal biology.

At a fundamental level, tissue clearing approaches render naturally opaque biological specimens optically clear. This transparency facilitates imaging large tissue volumes in their native, three-dimensional state using light microscopy approaches at imaging depths and resolutions that would otherwise not be possible due to scattering and diffusion of light ^14^.

Irrespective of the specific chemical reagents employed, all current tissue clearing techniques achieve transparency through similar physical principals. These approaches seek to minimize differences in refractive indices (RI) throughout a sample, as well as between the sample and the imaging media, thereby facilitating the passage of photons from a light source (excitation) through a tissue (emission) to ultimately reach a detector ^14,17^. The core principles of tissue clearing are removal of light-scattering lipids (RI∼1.47) and exchange of intracellular and extracellular fluids (RI∼1.35) for a solution with an RI equivalent to the protein and nucleic acid constituents that are left behind in the cell (RI>1.50), with the goal of creating a uniform density of scatterers so that all wavelengths of light pass through a tissue ^17^. As the underlying principles of tissue clearing have been described elsewhere in detail ^18^, and recent clearing approaches have been well summarized by others ^14^, we will only briefly discuss the logic underlying tissue clearing. At a gross level, tissue clearing techniques can be divided into either solvent-based (hydrophobic) or aqueous-based (hydrophilic) approaches. Early on, solvent-based methods were faster and generated a final RI more closely matched to that of the delipidated, protein-rich cleared sample. However, these methods also induced significant tissue shrinkage and generally eliminated signal from fluorescent reporters due to the requisite dehydration of the sample prior to delipidation ^19^. On the other hand, aqueous-based methods preserved signal from endogenous fluorescent proteins and maintained (or in some cases increased) sample size due to osmosis ^18^.

The conceptual divide that once viewed these two approaches as incompatible has been replaced with the understanding that steps from each methodology can, and should, be combined to optimize tissue transparency. As suggested by Richardson and colleagues, tissue clearing should be viewed as a pipeline consisting of the following modules: 1) sample fixation, 2) optional pre-treatment(s) (decolorization, decalcification, hydrogel embedding, dehydration), 3) delipidation (either active or passive), 4) optional fluorescent labeling (antibody, nanobody, dyes), 5) RI matching, and 6) image acquisition and analysis ^14^.

Early solvent- (i.e. BABB, 3DISCO), aqueous- (i.e., CUBIC, ClearT, ScaleAS, SeeDB) and aqueous hydrogel-based (CLARITY, PACT-PARS, SHIELD) clearing methods focused on RI matching and on lipid removal in the brain ^20^. However, these methods failed to clear thicker specimens, which blocked photons in the visible spectrum (400-600 nm) due to light scattering from mismatched RIs and light absorbance by endogenous chromophores such as hemoglobin and myoglobin ^21^. Attempts to decolorize tissues through either peroxide bleaching ^22^, or altering the pH of samples, attenuates signal from GFP-related fluorescent proteins ^23–25^. However, subsequent testing determined that amino alcohols could decolorize PFA-fixed by eluting heme ^26,27^. Overall, adapting clearing techniques for larger samples, particularly those containing muscle and mineralized tissues, such as bones or teeth, presents a formidable challenge.

While the composition of dense collagen fibers embedded within a highly calcified extracellular matrix, primarily made of hydroxyapatite crystals, imparts bone with mechanical strength, it also scatters light, preventing photons from passing through the tissue ^28,29^. Further, the extensive collagen fiber networks, minerals and proteoglycans of bone and muscle produce high autofluorescence upon fixation, potentially obscuring fluorescent signals of interest ^30–32^. In addition to these obstacles, the varying densities and mineral content of the honeycomb-like, porous network of plates and rods in trabecular bones versus the dense, solid cortical bones that resists decalcification and clearing can yield uneven clearing results within the same sample ^31^. Complicating these efforts, each of the tissues within the knee feature different refractive indices (RI), including bone (RI ∼ 1.50) (Rahlves et al., 2013), articular cartilage (RI ∼ 1.35) (Wang et al., 2010), and surrounding muscles (RI ∼ 1.38) (Deng et al., 2016). Thus, a prerequisite to such imaging studies is the development of a robust tissue clearing protocol that renders all components of the knee (or any complex joint) optically transparent, as many of these early clearing methods achieved only modest imaging depths of around 200 μm when applied to bone ^29^.

Aqueous, hydrogel-based methods, such as PACT (passive CLARITY) and PARS (perfusion-assisted agent release in situ), stabilize tissue during lipid extraction, but failed to achieve optical access beyond a depth of 200-300 μm. The CLARITY variant PACT-deCAL incorporated EDTA decalcification at 37°C *after* hydrogel embedding and SDS delipidation, although only staining for DRAQ5 dye was shown in this study ^33^. However, Bone CLARITY surpassed these limitations by adding 2 weeks of decalcification prior to hydrogel stabilization and SDS-mediated delipidation followed by amino alcohol (Quadrol) treatment to minimize heme autofluorescence, with all steps carried out under convective flow (via heat). RI matching in RIMS (RI=1.47) and LSFM imaging allowed them to visualize genetically labelled (*Sox9^CreER+^*; *R26^lsl-tdTomato+^*) fluorescent cells in the femur, tibia, and vertebral column at reported depths of up to 1.5 mm, with a high signal-to-noise ratio (SNR) ^34^. However, this method was only applied to isolated bones and not intact joints, although more recent attempts using hydrogel have also provided encouraging results.

In the hydrogel-based, passive clearing method Bone-mPACT+, Cho and colleagues decalcified bones in 11-13 days with one of four reagents (20% EDTA, Calci-Clear, 5% nitric acid, or 10% formic acid), which reduced the tissue damage associated with CLARITY-based methods by adding an additional detergent to the delipidation step. Furthermore, they also included α-thioglycerol to prevent tissue discoloration, then removed heme with 25% triethanolamine (TEA) (rather than Quadrol) before RI matching in *n*RIMS (RI=1.46), which yielded clear signal in *Cx3cr1^GFP^* mouse humerus and femur^30^. While CLARITY-based clearing may not yield the optical transparency achievable with clearing methods using organic solvents (e.g., BABB, 3DISCO, iDISCO, and PEGASOS) ^35^, the comprehensive comparison of bone samples across various methods suggests Bone-mPACT+ may be quite effective ^30^.

While these methods relied on genetically encoded fluorescent reporters, modified whole body clearing approaches have been adapted to image structures in the bones, such as the 3DISCO variant uDISCO, which adds Vitamin E to scavenge peroxides and the more stable *tert*-butanol for dehydration rather than THF, followed by delipidation with DCM and RI matching in a mixture of BABB and diphenyl ether (BABB-D), rather than DBE or BABB (both of which can form peroxidase and quench the fluorescence signal due to the presence of reactive benzylic C-H and C-O bonds). While uDISCO preserved GFP signal better than several other methods (CUBIC, SeeDB, ScaleS, 3DISCO, and PACT), it induced significant isotropic shrinkage of tissues and pigmented heme remained in the bones ^36^.

Notably, signal from endogenous fluorescent proteins may not overcome autofluorescence from skin, muscle and calcified bone. Accordingly, whole body clearing methods have been combined with immunostaining to enhance fluorescence signal. vDISCO (nanobody or V_H_H-boosted DISCO) employs whole-body perfusion mediated nanobody immunolabelling in conjunction with whole-body tissue clearing. After fixation, samples are decolorized using aminoalcohols (Quadrol), decalcified with EDTA, followed by nanobody perfusion and subsequent 3DISCO clearing (THF, DCM, BABB) and imaging. While vDISCO eliminates endogenous fluorophores, it enables robust nanobody labeling in the bone ^37,38^.

Continuing to expand these solvent-based approaches, Erturk and colleagues’ recent wildDISCO method combines perfusion-mediated fixation and decalcification with EDTA, followed by forced circulation of standard IgG antibodies in the presence of cyclodextrin and 3DISCO clearing (THF, DCM, BABB). The advantage of cyclodextrin is twofold, as it depletes cholesterol from membranes, while also reducing antibody aggregation, which identified proliferating Ki67^+^ cells in the bone marrow ^39^.

Finally, PEGASOS employed 5 days decalcification (20% EDTA) followed by decolorization with Quadrol and ammonium, delipidation through an increasing gradient of tert-butanol (uDISCO) in the presence of Quadrol (which raised the pH to over 9.5), dehydration in tert-butanol/polyethylene glycol and RI-matching in benzyl benzoate/PEG/quadrol (RI=1.54).

Despite these efforts, further optimization is needed to reduce autofluorescence from the surrounding muscle and bone and enable high-resolution imaging of bone and the surrounding tissues.

Neurovascular networks within the joint are thought to play critical roles in pain perception, inflammation, and tissue remodeling in osteoarthritis ^40^. Motivated by our interest in mapping vascular and nervous system innervation in health and disease ^41^ and the fact that neurovascular remodeling due to aging and disease condition is not yet fully explored in 3D, hence there is need for a robust clearing method that will allow for neurovascular imaging of joint tissue *in situ*.

Our results demonstrate that extended decalcification and delipidation in a modified iDISCO^+^-based clearing protocol most effectively render the murine hindlimb, and thus the knee joint, optically transparent. We also emphasize the critical role of refractive index-matching media in achieving imaging clarity and provide recommendations for adapting protocols to aged and pathological bone samples. This work provides a refined framework for studying neurovascular and structural changes in bone, with implications for understanding the pathological progression of diseases such as osteoarthritis.

## MATERIALS AND METHODS

### Animal Husbandry

All mouse protocols were approved by the Institutional Animal Care and Use Committee (IACUC) at the University of Virginia School of Medicine. For all experiments, the day of birth was considered P0, and all adult mice were 8 weeks of age (or older, as noted in the text). Mice were housed with access to food (normal chow diet) and water ad libitum on a 12-h light–12-h dark cycle at 21 °C and 50–60% humidity.

### Lectin Perfusion and Vascular Labelling

Mice were prepared for vascular labelling as previously described in detail ^42^. Briefly, mice were transferred to a secure induction chamber (approximately 1L in volume for an adult mouse) and anesthetized using 4.0% isoflurane (1-2 L/min flow rate). Once the animal failed to maintain its righting reflex, and breathing had slowed, they were removed from the induction chamber and placed on their back while a tight-fitting cone was placed over their nose, and anesthesia maintained using 2-3% isoflurane. After the depth of anesthesia was confirmed by absence of toe pinch reflex, mice were retro-orbitally injected with 50 μL *Lycopersicon esculentum* (tomato) lectin 649 nm (Vector Laboratories, USA DL-1178-1) into the retro-bulbar sinus vein using a 31-gauge needle. The needle was then gently removed, and lectin was allowed to circulate for 15 minutes. The animal was then transferred to a Styrofoam board, their extremities pinned, the chest sprayed with 70% ethanol, and the dermis, rib cage, and then peritoneum opened to allow access to the heart. Then 100-150 µL of lectin-649 nm was perfused transcardially through the left ventricle using a 27-gauge needle, the lectin was allowed to circulate for 5 minutes with the needle in place at the site of injection. Next, the mice were perfused by 10 mL of 1x PBS, then 10 mL of ice-cold 4% PFA/1x PBS with the help of syringe pumps (InfusionONE NE-300, New Era Instruments, USA) set at 0.4 mL/ minute with the right atrium punctured to allow for venous blood draining out and efficient perfusion.

### Hindlimb Preparation and Decalcification

Following successful perfusion, hindlimbs were dissected away from the mouse, de-skinned, and submerged in 50 mL of 4% PFA/1x PBS in a 50 mL falcon tube (VWR 10025-682), and kept on an orbital shaker set at 30 rpm, protected from light, overnight at 4°C. The following day, limbs were washed 3 times in 1x PBS for 30 minutes each at room temperature on an orbital shaker set at 30 rpm. The hindlimbs were then decalcified in 50 mL of 10% EDTA/1x PBS in a 50 mL falcon tube at room temperature on an orbital shaker set at 30 rpm for either 2 or 5 days (depending on the age of the sample). Samples were then washed 3 times with 1x PBS for 30 minutes each at room temperature on an orbital shaker set at 30rpm and processed for tissue clearing as indicated below.

### iDISCO^+^ Tissue Clearing

Samples were processed as described^43^. Fixed, decalcified hindlimbs in 50 mL falcon tubes (VWR 10025-682) were serially dehydrated by 60-minute washes through an increasing gradient (20%, 40%, 60%, 80%, 100%) of methanol (Sigma-Aldrich 154903) diluted in water with gentle agitation, while protected from light, followed by overnight incubation in 100% methanol. Hindlimbs were then delipidated in 66% v/v dichloromethane (DCM) (Sigma-Aldrich 270997) prepared in methanol for 24 hours at room temperature on an orbital shaker. The following day, the samples were washed two times, for 15 minutes each wash, in 100% DCM to remove methanol. The samples were then incubated for 24 hours at room temperature, with gentle agitation, in 100% dibenzyl ether (DBE) (ThermoScientific, A18447.30) to render the samples transparent. The following day, the DBE was exchanged for fresh solution and samples were placed on an orbital shaker at room temperature, protected from light, for another 24 hours. Samples were then stored in DBE at room temperature until they were imaged.

### wildDISCO

Samples were processed as described^39^. Briefly, samples were placed in a 20 mL glass scintillation tube (Sigma-Aldrich DWK986546) wrapped in foil to protect them from light. Samples were delipidated by a series of 12-hour washes through an increasing gradient (50%, 70%, 80%, 100%) of tetrahydrofuran (THF) (Sigma-Aldrich 186562) diluted in water with gentle agitation. Next, samples were washed in 100% DCM for 3 hours at room temperature on an orbital shaker with gentle shaking. Samples were then rendered transparent in BABB, a mixture of Benzyl alcohol (Sigma-Aldrich24122) and benzyl benzoate (Sigma-Aldrich W213802) at a ratio 1:2, respectively, for 24 hours with gentle agitation in the same scintillation tube. Samples were stored in a fresh BABB solution at room temperature until they were imaged.

### Binaree Tissue Clearing

Hindlimbs were processed using the Binaree Rapid Tissue Clearing system (Binaree, Inc., BDTC-003), as recommended by the manufacturer. Briefly, fixed, decalcified hindlegs were placed in the sample chamber and a sponge was placed on top of the sample to keep it in place. Next, the sample was submerged in 20 mL of Rapid Binaree Tissue Clearing Rapid Solution (Binaree, BRTC) that was pre-equilibrated to 37°C. The chamber was secured to the electrode, and the machine set to 100 V for 24 hours for electrophoretic clearing. The following day, the sample was washed in ddH_2_O three times, for 10 minutes each wash, in a 20 mL glass scintillation tube wrapped in foil to protect it from light. The samples were then submerged in EZ View solution [80% Nycodenz (PROGEN 18003)/7 M Urea (Sigma-Aldrich 51456)/0.05% sodium azide (Sigma-Aldrich S2002)/0.02 M sodium phosphate buffer] for 24 hours with gentle agitation at room temperature to render the samples transparent, while still protected from light. The EZ view solution was replaced the following morning and samples were stored in EZ view at room temperature until imaging.

### CLARITY Tissue Clearing

X-CLARITY clearing was performed according to the manufacturer’s instruction (Logos Biosystems). Briefly, after perfusion and fixation, mouse hindlimbs were immersed in 20 mL of X-CLARITY Hydrogel Solution with 0.25% (w/v) of polymerization initiator VA-044 (Logos Biosystems, C1310X) and incubated at 4°C for 24 hr. Cross-linking of the hydrogel was then induced thermally by incubating at 37°C under vacuum (–90 kPa) for 3 hr. After the crosslinking reaction, the hydrogel solution was removed and the hindlimbs were washed three times in 1x PBS, for 1 hr each wash, followed by an additional overnight wash at 4°C. The hydrogel infused and crosslinked hindlimbs were then submerged in electrophoretic tissue clearing solution (Logos Biosystems, C13001) and cleared using the X-CLARITY Tissue Clearing System (Logos Biosystems) with the following settings: 1.0 A and 37°C for 24 hr. After electrophoresis, hindlimbs were washed three times in 50 mL of 1x PBS, 1 hr each, then once overnight at room temperature. Samples were then RI matched in EZ View solution for 24 hours on an orbital shaker, and stored in fresh EZ view solution until they were imaged.

### EZ Clear Tissue Clearing

Samples were cleared as described previously^42,44^. Briefly, fixed, decalcified samples were submerged in 50%, 70%, then 80% Tetrahydrofuran (THF) (Sigma-Aldrich 186562-1L) diluted in ddH_2_O inside of a 20 mL glass scintillation tube (Sigma-Aldrich DWK986546) with the exterior wrapped in foil to protect the samples from light, and then incubated for 12 hours in each solution on an orbital shaker set at 100 rpm at room temperature. Samples were then washed four times in ddH_2_O, for 60 minutes each wash, to remove the THF solution. The vial was left open after the last wash to allow for evaporation of ddH_2_O. Next, the samples were submerged in EZ View solution in same scintillation tube wrapped in foil to protect the sample from light. Samples were stored in a fresh EZ View solution in the 20 mL glass scintillation tube until imaging.

### Vascupaint Perfusion and Micro-CT Imaging

Adult C57BL/6 male and female mice were anesthetized using vaporized isoflurane until they were unresponsive to noxious stimuli. Afterward, their chest was opened, rib cage reflected, and right atrium opened. Then, the animal was transcardially perfused through the left ventricle using a blunted 25-gauge syringe (BD PrecisionGlide, #305122) with 8 mL warm 1x PBS / 20 U/mL heparin (Mckesson, #63739092025), followed by 8 mL 10% neutral buffered formalin (Leica, #3800598), and then 1 mL Vascupaint^TM^ (MediLumine Inc, MDL-121). Vascupaint^TM^ was prepared as follows: 1 mL silicone, 2 mL diluent, and 40 μL catalyst. The following day, the leg was removed from the spine near the hip joint, and then the skin and fur were removed. Hindlimbs were then embedded in 1% low melt agarose in ddH_2_O in a 5 mL plastic vial (Axygen Scientific, ST-5ML) to prevent sample movement during imaging and to prevent tissue shrinkage due to dehydration. Micro-CT images were obtained using a SkyScan 1276 CMOS EDITION (at the Molecular Imaging Core at the University of Virginia School of Medicine). Scans were performed using the following parameters: source voltage and current (60 kV and 80 μA) and an exposure time of 471 milliseconds with an angular rotation step of 0.3° and an imaging voxel size of 9 µm^3^ and an AI filter of 0.25 µm. The distance of X-ray to object was set to 199.98 mm. Acquired images were then reconstructed using NRecon Reconstruction software (Bruker Micro CT, Kontich, Belgium). Micro-CT Tiff files were loaded on the NRecon software, after which the preview setting was selected. Background noise was reduced using the slider on the histogram on the output window to reduce background noise and ring artefacts, the ‘HU’ and ‘Scale’ tab were unchecked, but the ROI is selected. The new set of ‘tiff’ files were then saved in a separate folder. The saved NRecon tiff files were loaded into Imaris software V10.2.0 (Oxford Instrument UK) by using the import file series function on Imaris to create a 3D volume image of the micro-CT data. Next, the bone was masked by using the surface creation function on Imaris using the machine learning algorithm to cover the bone signals, the bone signal was masked by setting the voxel within the surface to zero and un-checking the box of for determining voxel size outside the surface. The vessels were then traced using the filament tracing function on Imaris software to trace the vessels and the tracing was statistically color-coded based on vessel diameter for visualization. The snapshot of the bone surface with the statistically colour-coded vessel tracing were captured together on Imaris by using the snapshot function. The images were then exported to Adobe Illustrator for figure creations. Vascular tracing data were obtained on Imaris by clicking on statistics tab on the filament tracing window in Imaris software. Moreso, the filter function on the filament tracing was used to obtained data for different vessel diameter by adding the segment mean diameter filter to the result and different diameter set to obtain the data.

### Light Sheet Image Acquisition

Cleared mouse hindlimbs were imaged on a Cleared Tissue Light Sheet XL microscope (CTLS XL) (3i Intelligent Imaging Innovations, USA). The samples were mounted on the sample holder secured by a screw. The sample holder was then attached to the stage anchor inside the imaging chamber of the microscope secured by a magnet. The samples were placed in a way that they were submerged in a glass boat containing the respective refractive index matching solution. Images were captured using a 640 nm laser, set to 200 mW, with an exposure time of 100 ms. The images were captured at a resolution of 0.8 µm and depth of view of 11.0 µm in a tiled sequence with 15% overlap.

### LSFM Image Processing

Acquired tiled images were stitched together and merged into a 3D image using SlideBook Software (Intelligent Imaging). Merged images were converted to Imaris compatible files (.ims) using the ImarisFileConverter V10.1.0. (Oxford Instruments, UK). Images were then visualized and further processed using Imaris software V10.2.0 (Oxford Instruments, UK). Representative snapshots of maximum intensity projections (MIP) of the images were obtained using the snapshot function in Imaris, and snapshots of optical sections along the Z-axis were created using the oblique slicer function. For vascular analysis, the surface function in Imaris was used to trace vessels using the machine learning algorithm, then the surface tracing was masked by changing the voxel size within the surface to 200 and changing outside the surface to zero to produce a masked image of the vessels. Next, the filament tracing function in Imaris was used to color-code the vessels based on their diameter. All snap shot images were exported to Adobe Illustrator for creating figures. Mean fluorescence intensity of optical sections along the Z-axis were calculated using ImageJ software^45^.

### Statistical Analysis

Statistical analysis of analysis of variance (ANOVA), t-test, and multiple comparisons tests were performed using Prism 9 software (GraphPad).

## RESULTS

### iDISCO^+^ and EZ Clear yield superior signal to noise ratios in mineralized tissues

To visualize the complex network of vessels of the murine adult hindlimb, and specifically the vasculature surrounding and permeating the knee joint space, anesthetized mice were intravenously perfused with a far-red fluorescently conjugated lectin that specifically labels the endothelium^45–47^ (**Figure 1A-C**). Hindlimbs were then collected, fixed, and decalcified for two days in 10% EDTA at room temperature followed by tissue clearing and 3D imaging (**Figure 1D-H**).

**Figure 1:**
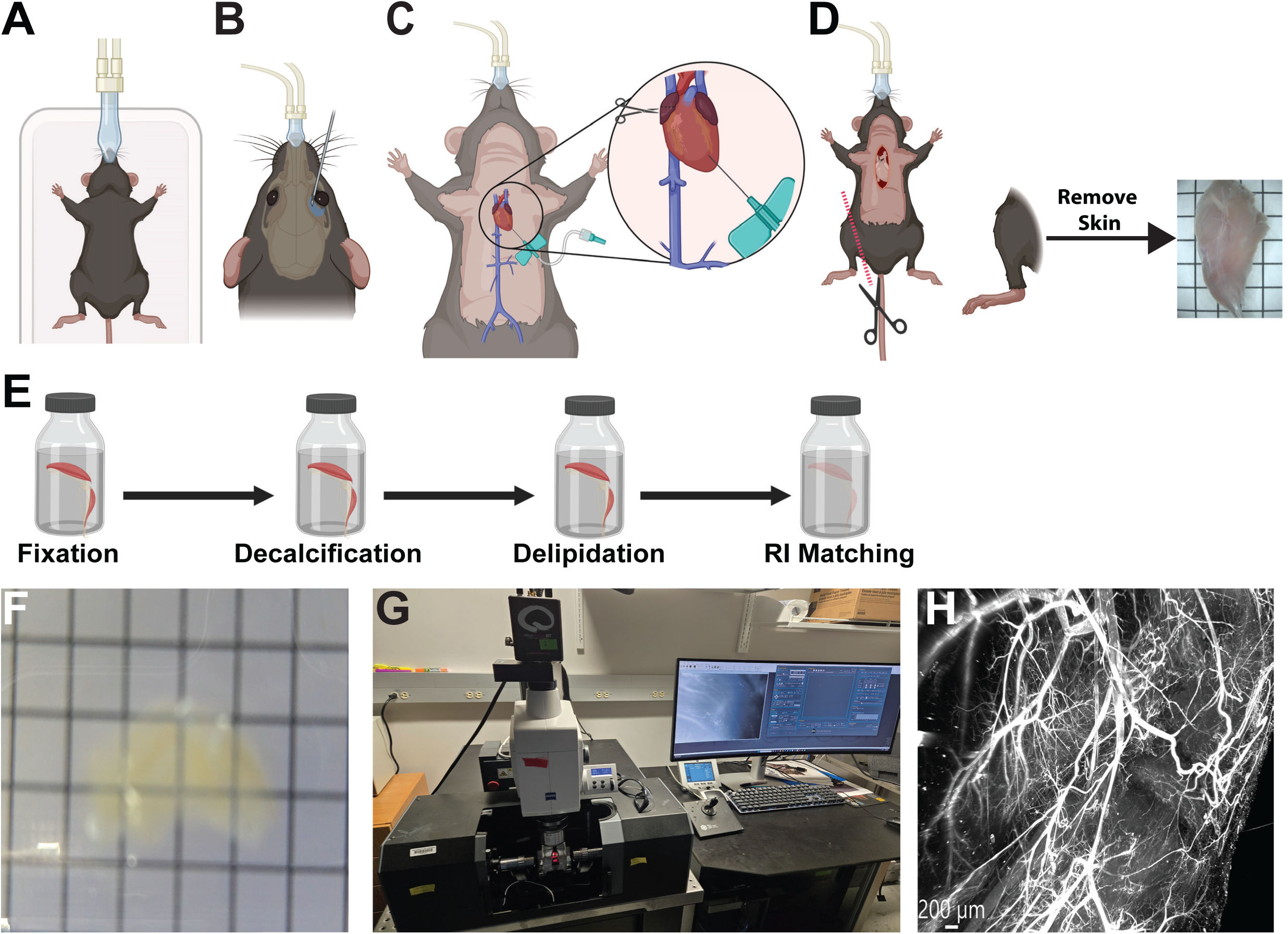
Overview of the experimental workflow. Animal is anaesthetized (**A**), followed by retro-orbital lectin injection (**B**). Animals then undergo transcardiac perfusion (**C**), followed by leg dissection and skin removal (**D**). (**E)** Leg samples undergo fixation, decalcification, delipidation methods and RI matching (tissue clearing) to render it transparent as shown in (**F**). Cleared leg samples were imaged on a Light-Sheet fluorescent microscope (**G**) to view the vasculature as shown in (**H**).

We performed a side-by-side comparison of several popular solvent-, aqueous-based, and hybrid tissue clearing methods to assess their efficacy at clearing mouse knee joints. The two solvent-based protocols we chose were variants of the original iDISCO approach^48^. Whereas iDISCO^+^ requires methanol dehydration and an overnight incubation in dicholoromethane (DCM) for delipidation followed by RI matching in Dibenzyl ether (BDE), wildDISCO delipidation uses tetrahydrofuran (THF) and a brief incubation in dichloromethane (DCM) followed by RI matching in BABB. We also included two commercial electrophoretic tissue clearing solutions: X-CLARITY (hydrogel-based) and Binaree Rapid Clearing. Finally, we also tested an updated variation of EZ Clear, a non-commercial aqueous approach we recently described, that relies on serial THF delipidation^42^, unlike the original protocol^44^. Following delipidation, all samples were then equilibrated in their respective RI matching media (**Figure 2**).

**Figure 2:**
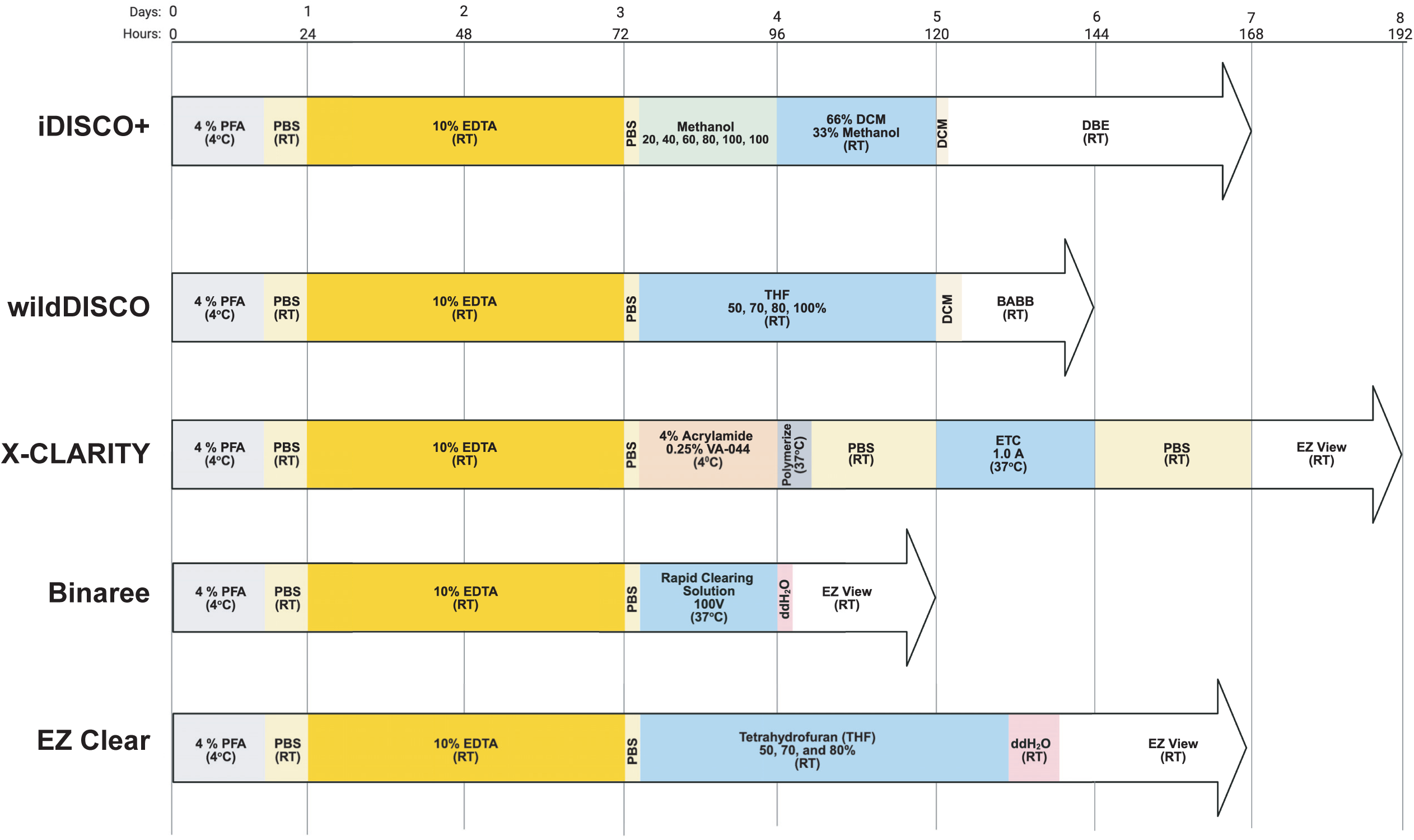
Comparison of the different solvent, hydrogel, and aqueous tissue clearing methods used in this study. A schematic showing the duration in time (hours and days), as well as the key steps and reagents, for iDISCO^+^, wildDISCO, X-CLARITY, Binaree and EZ Clear optical clearing methods.

Imaging by phase microscopy showed that the solvent-based tissue clearing methods wildDISCO and iDISCO^+^, along with the aqueous-based EZ Clear, effectively rendered the mouse hindlimb optically transparent. Notably, heme was still evident within the bones using these three methods, although less so in the wildDISCO processed samples (compare **Fig. 3B**, **C and F**). Neither the X-CLARITY or Binaree system achieved clearing comparable to iDISCO, wildDISCO, or EZClear (**Figure 3B-F**). Bone-CLARITY (which was not tested here) requires an extensive 2 week decalcification time^34^. Similar to our results, a recent comparative study of tissue clearing approaches showed that 3-4 days of decalcification with 10% EDTA followed by the original EZ Clear (a single incubation in 50% THF) protocol effectively cleared an isolated mouse tibia^30^.

**Figure 3:**
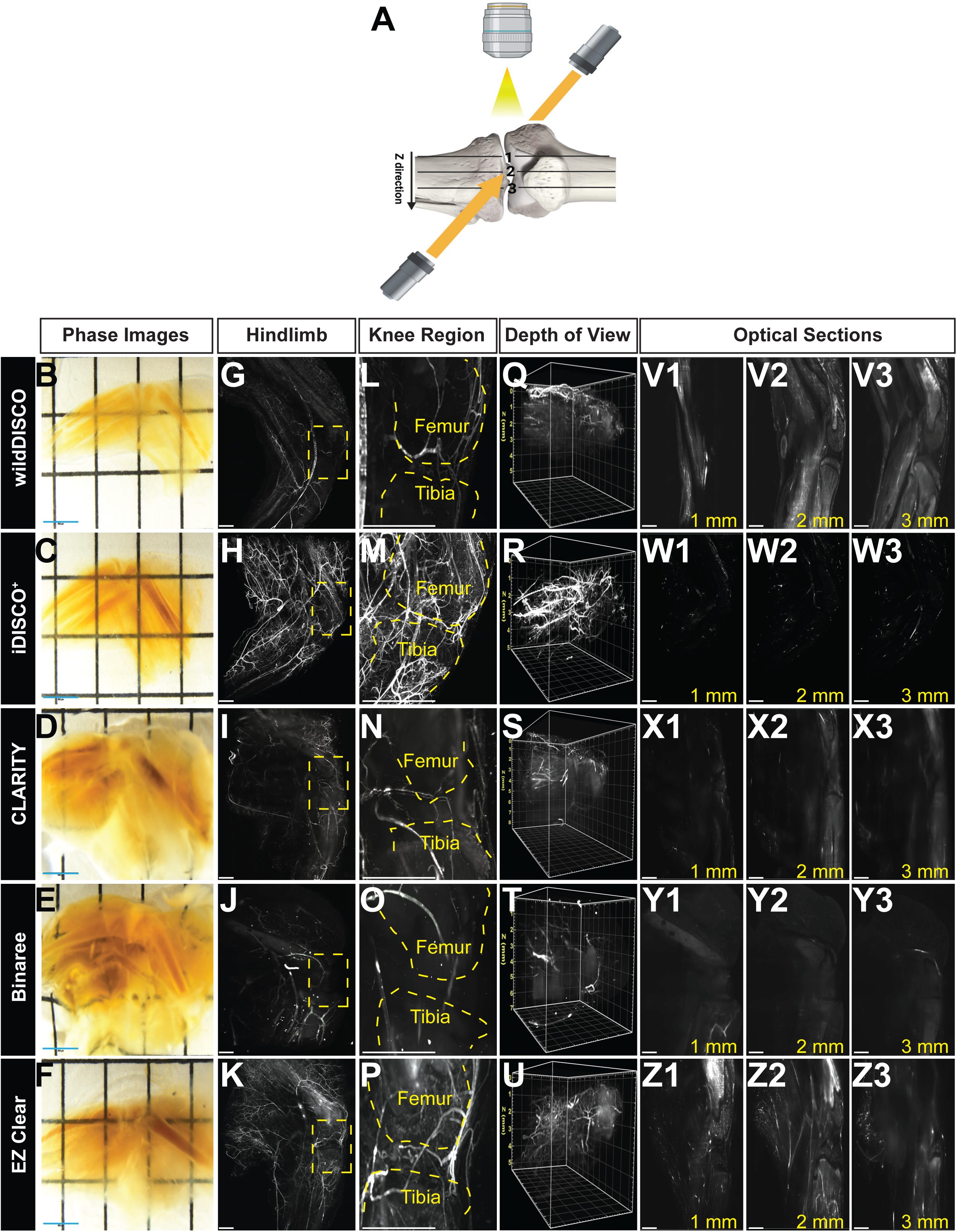
iDISCO^+^ optimally clears the mouse hindlimb. (**A**) Schematic diagram of the knee region indicating imaging orientation and planes of depth of view. (**B-F**) Phase images of mouse hindlimbs cleared using either wildDISCO, iDISCO^+^, X-CLARITY, Binaree, or EZ Clear. (**G-K**) Sagittal view of light sheet fluorescent microscope (LSFM) images of mouse hindlimbs following perfusion with lectin-649 nm and processing with the indicated tissue clearing protocols (far left column). Yellow dashed box indicates the knee region. (**L-P**) Magnified view of the knee region corresponding to the samples shown in panels G-K. (**Q-U**) Images showing the depth of view of the knee region (the yellow axis for the Z plane is indicated in each panel on the far left of the image). (**V** – **Z**) Optical sections along the Z-axis of the knee region at increasing depths (from 1 to 3 mm) highlight the retention of crisp signal in the vessels within the iDISCO+ and EZ Clear processed samples. Scale bars = 500 µm.

Following clearing, these samples were imaged by light sheet fluorescence microscopy (LSFM) with the laser beam directed at the leg sample from an antero-posterior plane while the medial side of the leg faced the objective lens (**Figure 3A**).

wildDISCO, iDISCO^+^ and EZ Clear facilitated photon travel through the samples, as fluorescent signal from the lectin dye revealed the details of the mouse hindlimb vasculature (**Figure 3G,H** and **K**). In contrast, X-CLARITY and Binaree processed tissues yielded less signal in the vasculature surrounding the mouse hindlimb, perhaps as expected given the phase microscopy results (**Figure 3I,J**).

In-depth examination of the tissues comprising the knee joint, which undergo extensive pathogenic remodeling in joint diseases such as osteoarthritis, revealed that iDISCO^+^, followed by EZ Clear, and to a lesser extent wildDISCO-processed samples, featured obvious vascular details and high fluorescent intensity for both superficial and deep vessels in and around the knee joint, with signal also evident in the femur and tibia. Conversely, in the knee joint space of X-CLARITY and Binaree cleared samples only peripheral vessels showed clear, sharp signal, with little fluorescence evident in either bone or the medial portion of the knee (**Figure 3L-P**).

To further characterize the efficiency of these clearing modalities, we evaluated photon penetration within the samples at various depths. Both solvent-based tissue clearing methods, iDISCO+ and wildDISCO, allowed photon penetration through the tissue, although wildDISCO samples showed decreased fluorescence intensity at greater depths (**Figure 3Q, R**). Binaree-and CLARITY-processed samples showed elevated background signal and decreased photon penetration (**Figure 3S,T**), while EZ Clear allowed deeper photon penetration throughout the tissue, with some loss of signal at greater depths (**Figure 3U**).

Virtual sections along the Z-axis (**Figure 3A**) revealed that lectin-based fluorescent signal was present in all planes for wildDISCO, iDISCO^+^, and EZ Clear processed samples, but diminished at deeper planes in Binaree and X-CLARITY treated hindlimbs (**Figure 3V-Y**). Quantification of signal to background ratio (SBR) on the LSFM images showed that iDISCO^+^ exhibited the greatest SBR of all five modalities (**Figure S1A**).

Consequentially, mouse hindlimb processed with iDISCO^+^ had the highest vascular mean intensity with the lowest background signal compared to other modalities (**Figure S1B**). Furthermore, quantification of mean fluorescent intensity of the optical Z-sections showed similar mean intensity in wildDISCO, iDISCO^+^ and EZ Clear processed samples which are significantly higher than Binaree and CLARITY modalities (**Figure S1C**). Collectively, these data show that iDISCO^+^, wildDISCO, and EZ Clear rendered mouse hindlimbs optically clear, enabling detailed LSFM imaging at depths of up to 5 mm, with iDISCO^+^ providing the most optimal results, followed by EZ Clear and wildDISCO.

### Aged mouse calcified tissues require increased decalcification

While many risk factors, including obesity, prior joint injuries, biological sex, and genetic disposition/congenital bone or ligament abnormalities correlate with the development of osteoarthritis, the most critical risk factor is age^49^. Accordingly, osteoarthritis disproportionately impacts the elderly^50,51^. The interplay between cellular senescence (particularly in articular chondrocytes and associated musculoskeletal cells), low-grade inflammation (especially in the synovium), and metabolic modifications synergize to drive the irreversible changes characteristic of age-related OA progression ^52–54^. Given iDISCO^+^ optimally cleared hindlimbs and knee joints of 2-month-old mice, we wondered if this same processing pipeline could effectively clear 6-month-old samples, an age at which mice have reached peak bone mass^55^. Surprisingly, while phase images suggested these aged tissues were of similar transparency as the 2-month-old samples (**Figure S2A, left panel**), LSFM revealed extensive background fluorescence signal (autofluorescence) in the femur, tibia, fibula, and patellar bones (**Figure 4A**). Higher-magnification imaging of the knee joint showed that this high background fluorescence partially obscured lectin-mediated fluorescence signal in the vessels around the knee joint of aged animals (**Figure 4B**), lowering the signal to noise ratio. Furthermore, while the depth of view indicated that photons were able to penetrate the samples, they appeared blurry at the thickest part of the aged sample (**Figure 4C**). Hypothesizing that the increased autofluorescence may be due to incomplete decalcification, the decalcification time of 6-month-old mouse samples was increased from 2 to 5 days prior to iDISCO^+^ clearing. LSFM imaging revealed a significant reduction in autofluorescence and increased signal to noise in the aged hindlimbs decalcified for 5-days (**Figure 4D**). Further examination of the knee region revealed detailed vessel structures with little background fluorescence signal evident from the bones upon extended decalcification (**Figure 4E**). Furthermore, photon penetration was increased in the aged samples that were decalcified for 5 days (**Figure 4F**). Examination of virtual sections at varying depths along the Z-axis revealed non-specific, background fluorescence at every depth examined in the 2-day decalcified tissues, particularly in deeper sections (**Figure 4G,H**). Conversely, the 5-day decalcified samples showed reduced autofluorescence in the bone and minimal non-specific background signal (**Figure 4I**). Quantification of lectin fluorescence indicates that 5-day decalcified samples featured higher signal and reduced background compared to 2-day decalcified samples (**Figure 4J, S2B**). Moreover, the mean fluorescent intensity across the Z-plane showed that 2-day decalcified samples had higher mean fluorescent intensity (**Figure S2C**) which is due to the high background fluorescence presence in the samples. Thus, imaging-based studies of aged mouse samples will likely benefit from increased decalcification prior to tissue clearing and imaging.

**Figure 4:**
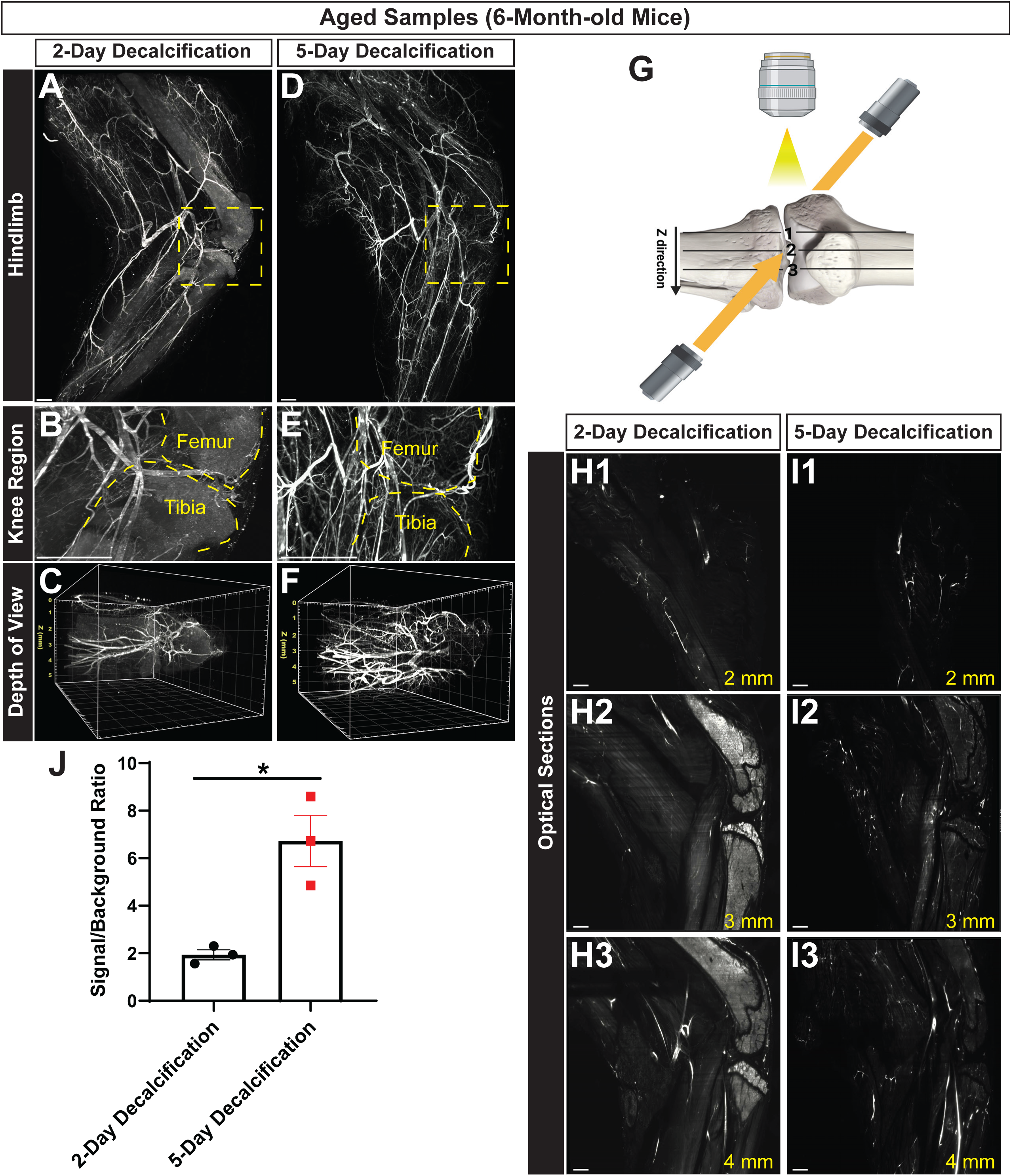
Aged samples require longer decalcification for optimal tissue clearing and imaging. (**A**) A sagittal maximum intensity projection following LSFM imaging of a mouse hindlimb perfused with lectin-649 nm and cleared using iDISCO^+^ with 2-days of decalcification in 10% EDTA. The yellow dashed area is magnified in (**B**) and represents the knee region, with the outline of the femur and tibia noted. (**C**) A depth of view image of the sample in panel A (note the z axis, in yellow, at the far left) showing how fluorescence signal diminishes at greater depths. (**D**) A similarly perfused mouse hindlimb processed for iDISCO^+^ clearing after 5-days of decalcification. (**E**) A magnified view of the knee region from panel A, (**F**) and a depth of view image showing improved signal intensity overall, less signal from bone, and more intense signal at greater imaging depths along the z axis. (**G**) Schematic of the knee region showing imaging orientation and planes of optical sections shown in panels H and I. (**H,I**) Comparison of optical sections of the knee along the Z-axis. (**J**) Quantification of the signal to background fluorescence ratio (SBR) in the mouse hindlimb showing increased SBR in the 5-day decalcification samples compared to 2-day decalcification (n=3 samples per treatment; t-test, p≤0.05). Scale bar = 500 µm.

### DBE refractive index matching media is superior to ECi and BABB for imaging iDISCO^+^ cleared knee samples

We next set out to identify the optimal refractive index (RI) matching media for iDISCO^+^-based LFSM imaging of musculoskeletal tissues that form the knee joint. Accordingly, we tested BABB (RI=1.559)^56^, Dibenzyl ether (DBE) (RI=1.562)^20^, and the non-toxic DBE alternative, ethyl cinnamate (ECi) (RI=1.558)^57^. Phase microscopy imaging showed that each of these three RI medias rendered iDISCO^+^ cleared mouse hindlimbs optically transparent to transmitted light, although more pigment was evident in the bones and blood vessels of the DBE-treated samples (**Figure 5A-C**). LSFM imaging confirmed that all three RI media preserved lectin fluorescence (**Figure 5D-F**). However, closer examination of the knee revealed that BABB-and ECi-matched samples were unable to resolve small diameter, deeper vessels were difficult to resolve and appeared blurry in BABB-and Eci-matched samples compared to those in DBE (**Figure 5G-I**). Furthermore, examining photon penetration indicated that BABB yielded aberrations and blurry signal (**Figure 5J**), as did ECi (**Figure 5K**), whereas DBE showed clear penetration throughout the tissue (**Figure 5L**). Optical sectioning revealed reduced lectin signal intensity in deeper planes of BABB and ECi-treated samples compared to those imaged in DBE (**Figure 5M-P**). Quantification confirmed that DBE had a significantly higher SBR than to either BABB or ECi (**Figure S3A**). Relatedly, while the mean fluorescence intensity within vessels was similar across all RI medias, DBE had the lowest background signal (**Figure S3B,C**). These data show that iDISCO^+^ cleared mineralized samples should be imaged in DBE for optimal penetration and superior signal to background noise intensity.

**Figure 5:**
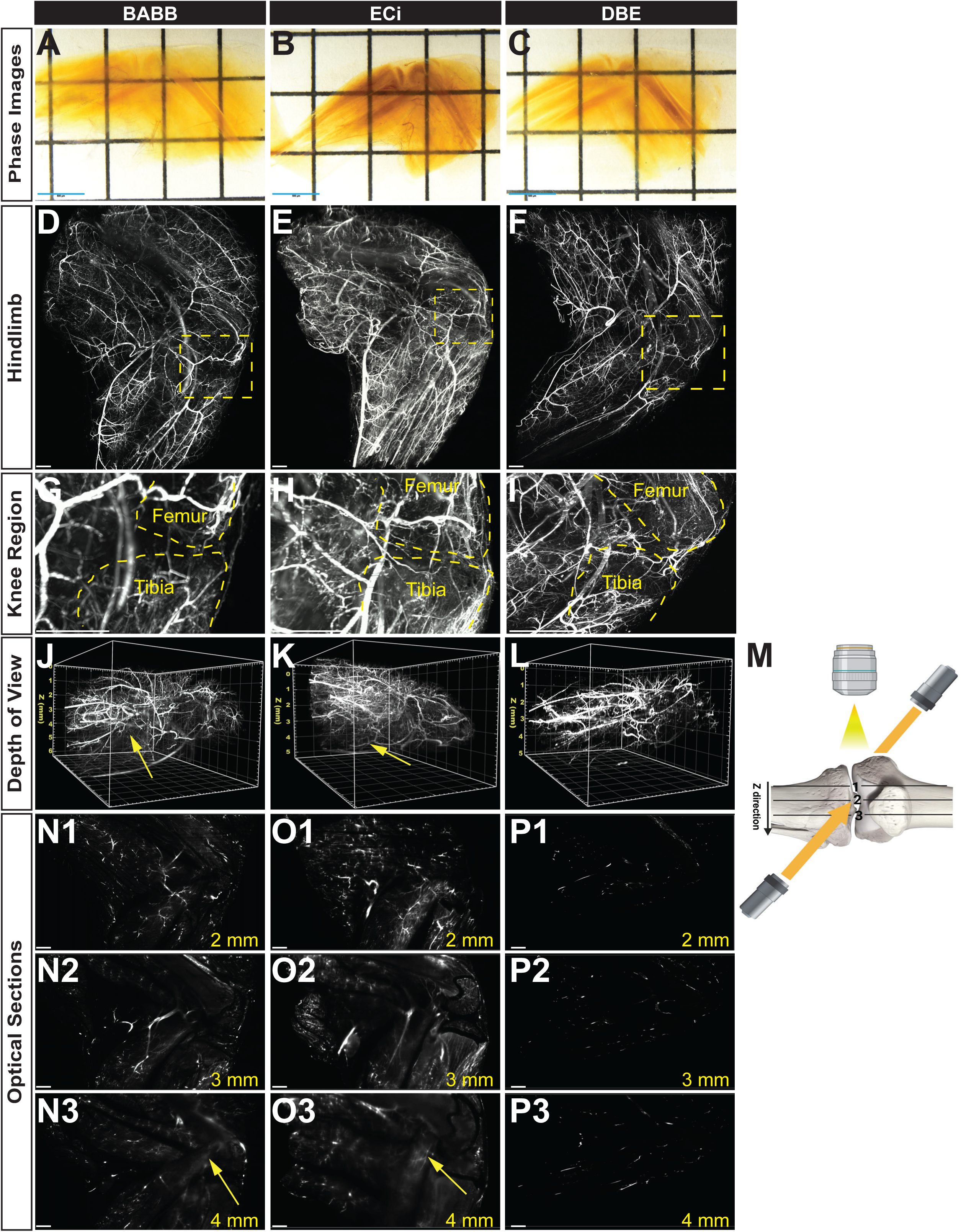
RI matching with DBE achieves better depth of view for iDISCO^+^ cleared mouse hindlimbs. (**A-C**) Phase images of mouse hindlimbs cleared with iDISCO^+^ and RI matched in either BABB, Ethyl Cinnamate (ECi) or Dibenzyl Ether (DBE). (**D**-**F**) Sagittal views of maximum intensity projections following LSFM imaging of mouse hindlimb RI-matched in different imaging medias. (**G-I**) Magnified views of the yellow boxed areas in D-F showing the knee region highlight the low background signal from the bone in DBE matched samples (unlike ECi). (**J** – **L**) Depth of view images show aberrations and background signal (yellow arrows) in the BABB and ECi samples. (**M**) A schematic diagram showing the imaging orientation and planes of optical sections shown in panels (**N** – **P**) along the Z-axis. Note the background from bone and muscle in the BABB and ECi samples. Scale bar = 500 µm

### iDISCO^+^ allows different image orientation acquisitions compared to EZ clear

Light propagation through biological samples is in part dependent on the distance photons must travel and the heterogeneity of the tissue being imaged^58^. As tissue thickness and imaging depth increase, so too does light scattering and absorption, which in turn reduce image clarity and signal intensity. To evaluate if the distance photons travel through cleared knee joints affects the quality of the resulting image, we tested if either imaging angle or sample orientation impact image quality using the best performing aqueous-(EZ clear) and solvent-based (iDISCO^+^) clearing methods. Images from an anterior orientation, where the laser beam is directed at the medial and lateral side of the hindlimb, while the collecting objective faced the anterior of the hindlimb, were compared to those obtained at a sagittal orientation, in which photons traveled from the antero-posterior plane and the objective faced the medial side of the hindlimb (**Figure 6A,B**). In the anterior orientation, superficial and peripheral vessels such as superior lateral and medial geniculate vessels (SLGV and SMGV) and inferior medial and lateral geniculate vessels (IMGV and ILGV), but not the deeper vessels of the hindlimb and knee region such as the popliteal artery (PA), were clearly identified in the EZ Clear sample (**Figure 6C**). Conversely, both peripheral and deep vascular structures of the hindlimb (SLGV, SMGV, ILGV, IMGV and PA) were evident in iDISCO^+^ samples (**Figure 6D**). However, imaging in the sagittal orientation improved resolution in both the superficial and deep tissues processed with EZ Clear (**Figure 6E, F**). Thus, decreasing the distance of photon travel through the sample improved overall imaging of the vasculature in the knee of EZ Clear processed samples (**Figure 6G,I**). However, both peripheral and deep blood vessels were well labelled following iDISCO^+^ processing, regardless of sample orientation (**Figure 6H,J**). Evaluation of photon penetration revealed decreased fluorescent signal at greater depths in the EZ Clear anteriorly oriented samples (**Figure 6K**,**M**). Photon depth was not impacted by sample orientation in the iDISCO^+^ cleared samples (**Figure 6L,N**). In agreement with these data, virtual sections demonstrated that EZ Clear samples imaged in the anterior orientation showed a depth-dependent decrease in signal intensity and diminished resolution of small vessels that directly correlated with depth along the z-axis (**Figure 6O**). Conversely, crisp, fluorescent signal from deep, small vessels was evident throughout the z-axis in iDISCO^+^ cleared samples imaged in the anterior orientation (**Figure 6P**). In the sagittal orientation, both methods successfully resolved small vessels throughout the depth of the sample (**Figures 6Q, R**). Quantification showed that the mean fluorescence intensity in EZ Clear samples is diminished in the anterior orientation compared to the sagittal orientation, whereas iDISCO^+^ shows a higher mean intensity in either view (**Figure S4A**). While both EZ Clear and iDISCO have similar lectin fluorescence signal within the vasculature, the anterior view of the EZ Clear had higher background than the sagittal view (**Figure S4B**) and the mean fluorescent intensity of the optical sections across the Z-plane showed that EZ Clear processed samples had higher mean fluorescence in both views than iDISCO^+^ due to background fluorescence (**Figure S4C**). These findings demonstrate that iDISCO^+^ provides superior sample clearing and imaging depth, regardless of imaging orientation, compared to EZ Clear.

**Figure 6:**
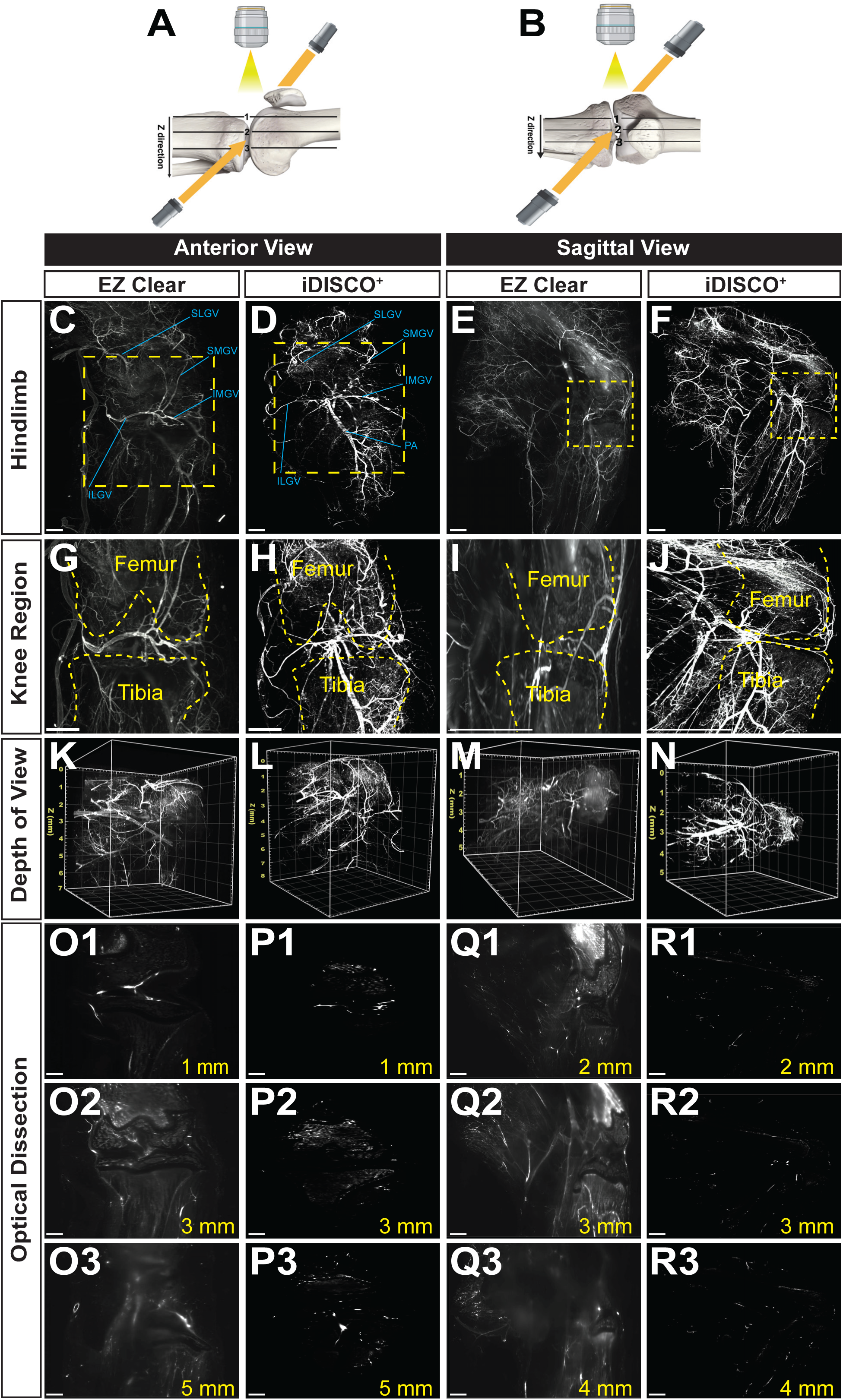
iDISCO^+^ tissue clearing improves fluorescence signal intensity and imaging depth in both anterior and sagittal imaging orientations. (**A**,**B**) Schematics illustrate the different imaging orientations and planes of optical sections for panels C-R. (**C-F**) Comparison of how an anterior or sagittal orientation of the sample relative to the microscope objective impacts fluorescence signal intensity and depth within the vasculature of the adult murine hindlimb following perfusion with lectin-649 and either EZ Clear or iDISCO^+^ tissue clearing. (**G-J**) Optical sections of both views, with the femur and tibia indicated. (**K**-**N**) Depth of view and (**O-R)** optical sections along the Z-axis of the knee region. Scale bar = 500 µm. SLGV = Superior Lateral Geniculate Vessel, SMGV = Superior Medial Geniculate Vessel, IMGV = Inferior Medial Geniculate Vessel ILGV = Inferior Lateral Geniculate Vessel.

### iDISCO^+^ -based clearing, followed by LSFM imaging of the vasculature in the mouse hindlimb outperforms micro-CT

Extensive studies of the mouse vasculature have been performed using micro-computed tomography (micro-CT) imaging^59–62^. Combined with a perfused contrast reagent to label the vasculature, this X-ray-based 3D-imaging modality, without the need for tissue clearing or for extensive immunolabeling methods, has been the gold-standard for capturing and analyzing vascular networks in mice^41,62,63^. However, micro-CT imaging has drawbacks^64^, as contrast agents that may alter biological structures, potentially interfering with signal from dense tissues (such as the bone) and masking details of interest (such as the microvasculature). Other potential confounds include exposure to potentially damaging X-rays (in the case of imaging living subjects), and the inability to resolve small caliber blood vessels due to the limited resolution of this method. To evaluate how iDISCO^+^-based tissue clearing and LSFM compares to contrast-assisted micro-CT, we compared the hindlimb vascular networks in animals processed for these distinct imaging modalities (**Figure 7**).

**Figure 7:**
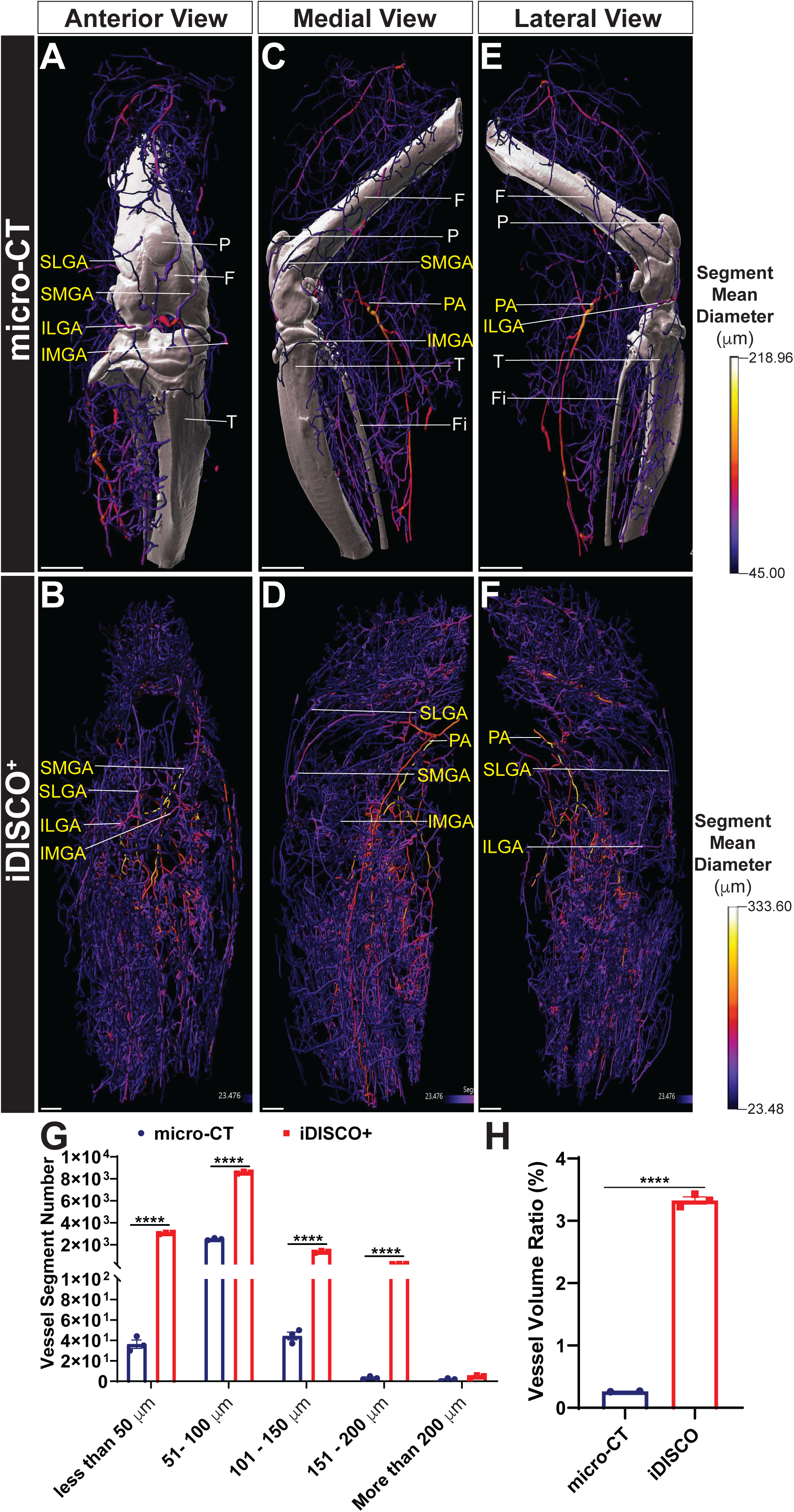
Lectin vascular labelling with iDISCO^+^-clearing reveals a more detailed view of the vascular within the mouse hindlimb than contrast-assisted micro-CT. (**A**,**B**) Anterior view of representative micro-CT images of the mouse hindlimb following perfusion with Vascupaint contrast agent and LSFM image of a mouse hindlimb perfused with lectin-649 and cleared using iDISCO^+^. Bone in the micro-CT images are pseudocolored white, while vessels in both the micro-CT and light sheet panels are color coded based on vessel diameter (the keys corresponding to vessel diameter are to the right of panels E and F). (**C, D**) Medial and (**E**,**F**) lateral views of the same samples. (**G**) Quantification of the frequency of different diameter vessels in micro-CT and LSFM imaged samples. (H) Quantification of the difference of vessel volume relative to the sample volume (calculated as Vessel Volume Ratio (%) = ^𝑉𝑒𝑠𝑠𝑒𝑙 𝑣𝑜𝑙𝑢𝑚𝑒^-_𝑆𝑎𝑚𝑝𝑙𝑒 𝑣𝑜𝑙𝑢𝑚𝑒_ x 100%) between micro-CT and LSFM imaged samples. F=Femur; Fi=Fibula; P=Patella; T=Tibia; IMGA=Inferior medial geniculate artery; ILGA=Inferior lateral geniculate artery; PA=Popliteal artery; SMGA=Superior medial genicular artery; SLGA=Superior lateral genicular artery).; Scale bar = 500 µm.

Large and medium sized arterial vessels were easily identified in both the micro-CT and LSFM-imaged samples, as the superior medial and lateral genicular arteries (SMGA, SLGA) and inferior medial and lateral genicular arteries (IMGA, ILGA), which together form the genicular anastomosis that supplies the knee region, as well as the popliteal artery, were all evident (**Figure 7**). 3D morphometric analysis revealed that micro-CT primarily captured arterial vessels larger than 45 µm in diameter, whereas vessels around 20 µm in diameter were readily evident in LSFM samples (**Figure 7G**). The overall number of vessels was greater in iDISCO^+^ imaged samples (**Figure 7H**). This is not unexpected, given the inability of Vascupaint to cross the small diameter (5 μm) capillary bed and enter the venous vasculature. Altogether, the ratio of vascular volume compared to total image volume was significantly higher in the lectin-labeled LSFM sample (**Figure 7H**). Together, these data demonstrated that tissue clearing followed by LSFM imaging enables visualization of a more extensive vascular network permeating the entire mouse hindlimb than micro-CT-based imaging.

## DISCUSSSION

Studying the mouse hindlimb and knee joint has been incredibly challenging due to the distinct optical properties of the heterogeneous tissues that comprise the leg, including muscles, tendons, ligaments, and bones. Traditional histological approaches, such as hematoxylin and eosin staining of 2D paraffin sections, have provided limited insights into this complicated joint system^65^. Even advanced modalities, such as immunostaining combined with confocal imaging of cryosectioned tissue, which is both laborious and time consuming, are often challenging to interpret. Overall, these 2D imaging techniques restrict analysis to a narrow tissue region within the microscope’s field of view, failing to capture the 3D architecture of the hindlimb in general, and knee in particular. As a result, spatial relationships of different tissues within the joint cannot be fully appreciated. To address these limitations, some researchers have explored 3D imaging modalities, such as micro-CT^60^. While this approach generates volumetric data, it suffers from limited resolution, and imaging at the cellular level is currently impossible using this modality. Furthermore, high-density bone tissue generates substantial background signals, often masking structures surrounding or embedded within the bone^66,67^.

Tissue clearing methods have emerged as a promising solution to these obstacles, rendering tissue optically transparent through lipid removal (delipidation) and refractive index (RI) matching to reduce light scattering^14^. These techniques have been successfully applied to visualize vascular and neural networks in the brain, as well as other organs, such as the kidney and liver ^44,68,69^. More recently, adaptations of tissue clearing protocols have extended their utility to bony structures^30,31,34^, indicating their potential for detailed analysis of the mouse hindlimb.

In this study, we systematically compared 2 organic solvent-based based methods (wildDISCO and iDISCO^+^), two commercially available electrophoretic approaches (CLARITY and Binaree), and our previously established aqueous-based technique, EZ Clear. We also investigated how sample age, RI media, and sample orientation influence imaging results.

Our results demonstrate that iDISCO^+^ and EZ clear enable high-resolution 3D visualization of fluorescently-labelled murine hindlimb vasculature. Notably, iDISCO^+^ allowed for deeper light penetration than all other methods tested. This indicates that the delipidation compound DCM used in iDISCO^+^, along with THF used in the EZ Clear approach, effectively remove lipids in the mouse hindlimb, allowing for effective tissue clearing, as previously reported^20,48,68^. We further observed that aged biological samples, which feature increased bone mineralization^70^, exhibited pronounced autofluorescence, which impaired image quality. This issue was mitigated by increasing the decalcification time, emphasizing the need for age and tissue specific optimization of any tissue clearing pipeline.

Another key determinant of clearing success is the choice of RI matching media. We compared several iDISCO^+-^compatible RI medias^39,57,69^ and found that DBE consistently yielded the best imaging results, with enhanced photon depth and a greater signal to noise ratio compared to BABB and ECi.

Importantly, we also found that – depending on the tissue clearing modality – sample orientation during imaging significantly impacts downstream imaging results. While the sagittal orientation enabled high quality imaging for both EZ Clear and iDISCO^+^, anterior-to-posterior imaging was only effective for iDISCO^+^-cleared samples. We attribute this difference to the anisotropic clearing efficiency of EZ Clear, particularly in thicker samples where complete delipidation and reaching RI equilibrium may be more difficult^42^. This result underscores the importance of optimizing sample orientation for each clearing protocol to ensure consistent and comprehensive visualization of structures of interest.

Comparison of tissue-cleared LSFM samples to micro-CT determined that clearing and LSFM provides superior resolution of small and medium caliber vessels that are typically not resolved using micro-CT due to limited contrast agent perfusion and signal interference from surrounding bone. Comparison to a blood-pool contrast agent, which could circulate through the entire vascular network (and not just the arterial endothelium), may provide a more appropriate comparison to lectin and LSFM imaging. However, in our experience these blood-pooling contrast agents induce significantly less signal attenuation than compounds such as Vascupaint, Microfil, or barium-based agents^62^.

Overall, these findings show the promise of iDISCO^+^ as a powerful tool not only for imaging the hindlimb vasculature, but also for investigating musculoskeletal pathologies such as ischemia, vascular calcification, inflammation, and age-related tissue degeneration.

Finally, our findings highlight the possibilities for integrating antibody-based immunostaining with tissue clearing to analyze region- and tissue-specific molecular changes in the knee. While immunostaining combined with clearing has been applied in other organs^39,48,69,71^, and even the femur^72,73^, few if any studies have demonstrated its effectiveness in intact musculoskeletal joints. Further optimization of antibody penetration, staining conditions, and decalcification protocols will be essential for enabling cell-type-specific and pathological studies of this complex anatomical region.

## ACKNOWLEDGEMENTS

The authors thank members of the RE-JOIN consortium.

## AUTHOR CONTRIBUTIONS

JDW conceptualized the study. JDW and AIO wrote the original draft. AIO, AP, TA, and CWH executed, imaged, and analyzed experiments. AIO, TA, CWH, and JDW, were involved in the design of experiments. NH revised the manuscript and provided helpful feedback and comments. All authors edited the manuscript and consented to its contents.

## SUPPLEMENTARY FIGURE LEGENDS

**Figure S1:**
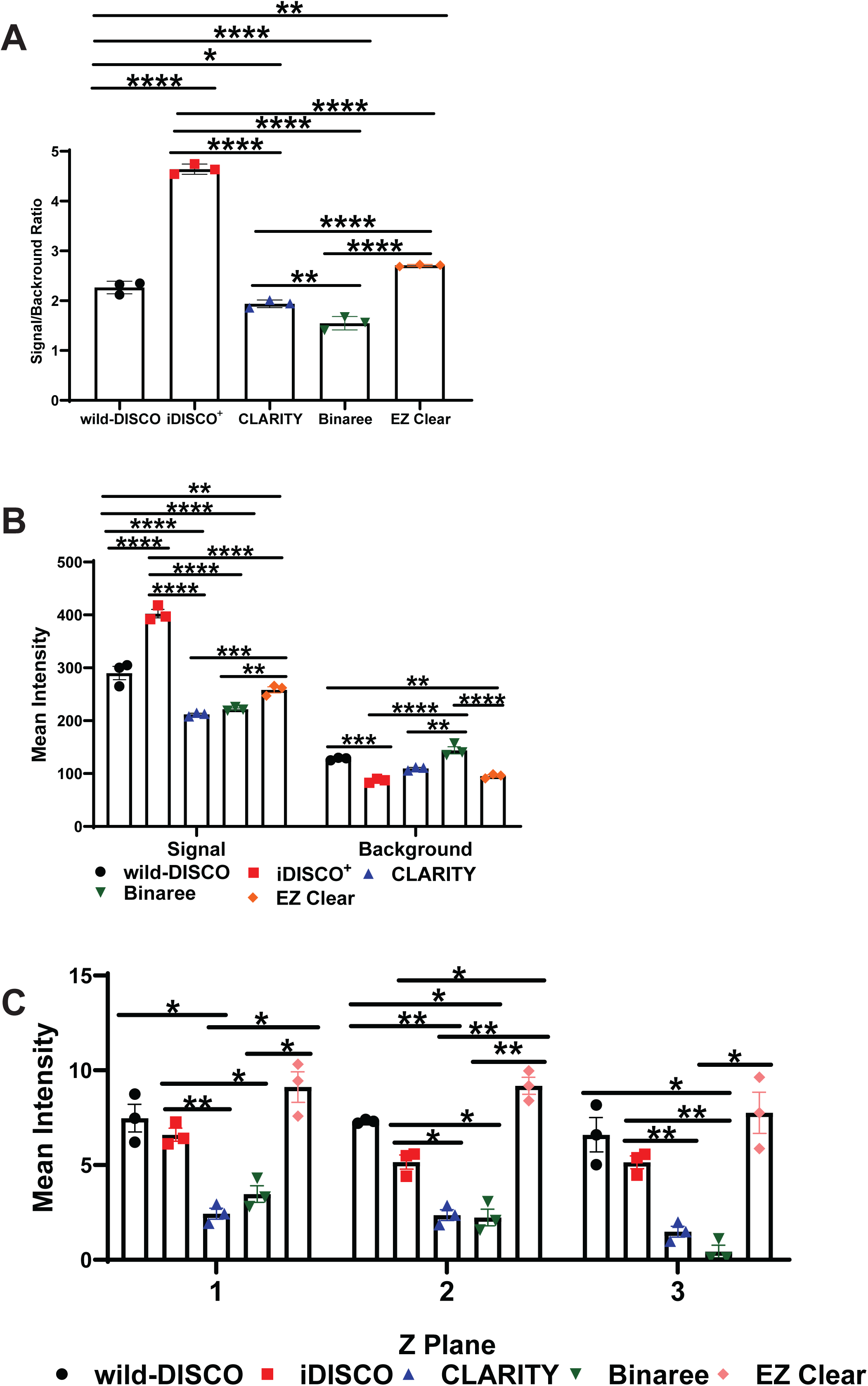
Comparison of the efficacy of different tissue clearing modalities on mouse hindlimb labelled with lectin-649. (**A**) Quantification of signal to background fluorescence ratio (SBR). (**B**) Quantification of the mean intensity in lectin labelled vessels (i.e. “signal”) and bone autofluorescence (i.e. “background”). (**C**) Quantification of mean fluorescent intensity in the optical sections across the Z-plane.

**Figure S2:**
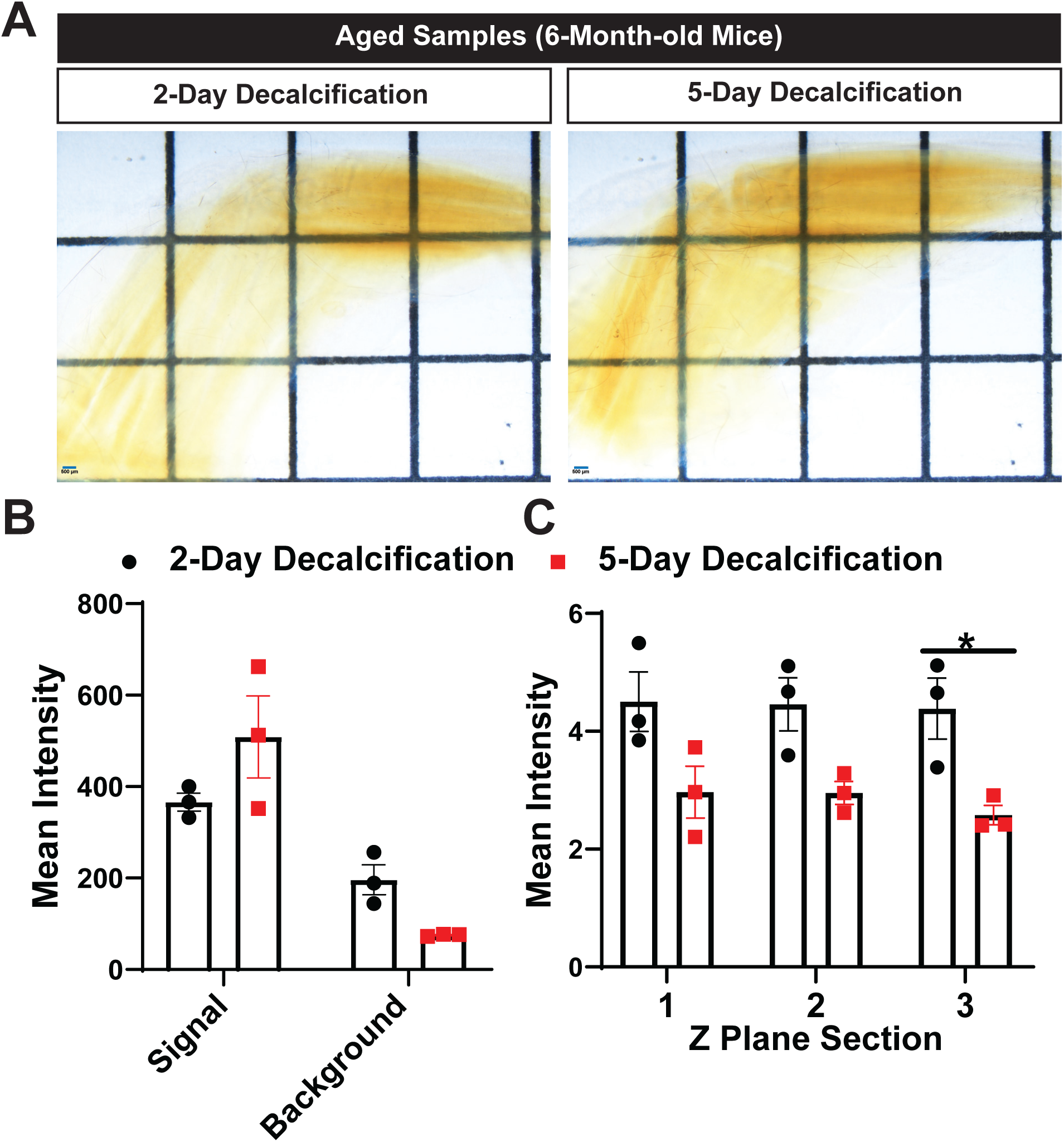
Impact of decalcifying time on clearing results of aged (6-month-old) mouse hindlimbs. (**A**) Phase images of hindlimbs from 6-month-old mice cleared with iDISCO^+^ after 2- or 5-days of decalcification in 10% EDTA. (**B**) Quantification of the mean fluorescence intensity of lectin labelled vessels (i.e. “signal”) and bone autofluorescence (i.e. “background”) for both treatment regimens. (**C**) Quantification of mean fluorescence intensity in optical sections along the Z-plane (increasing depths from sections 1 to 3). p ≤0.05, t-test.

**Figure S3:**
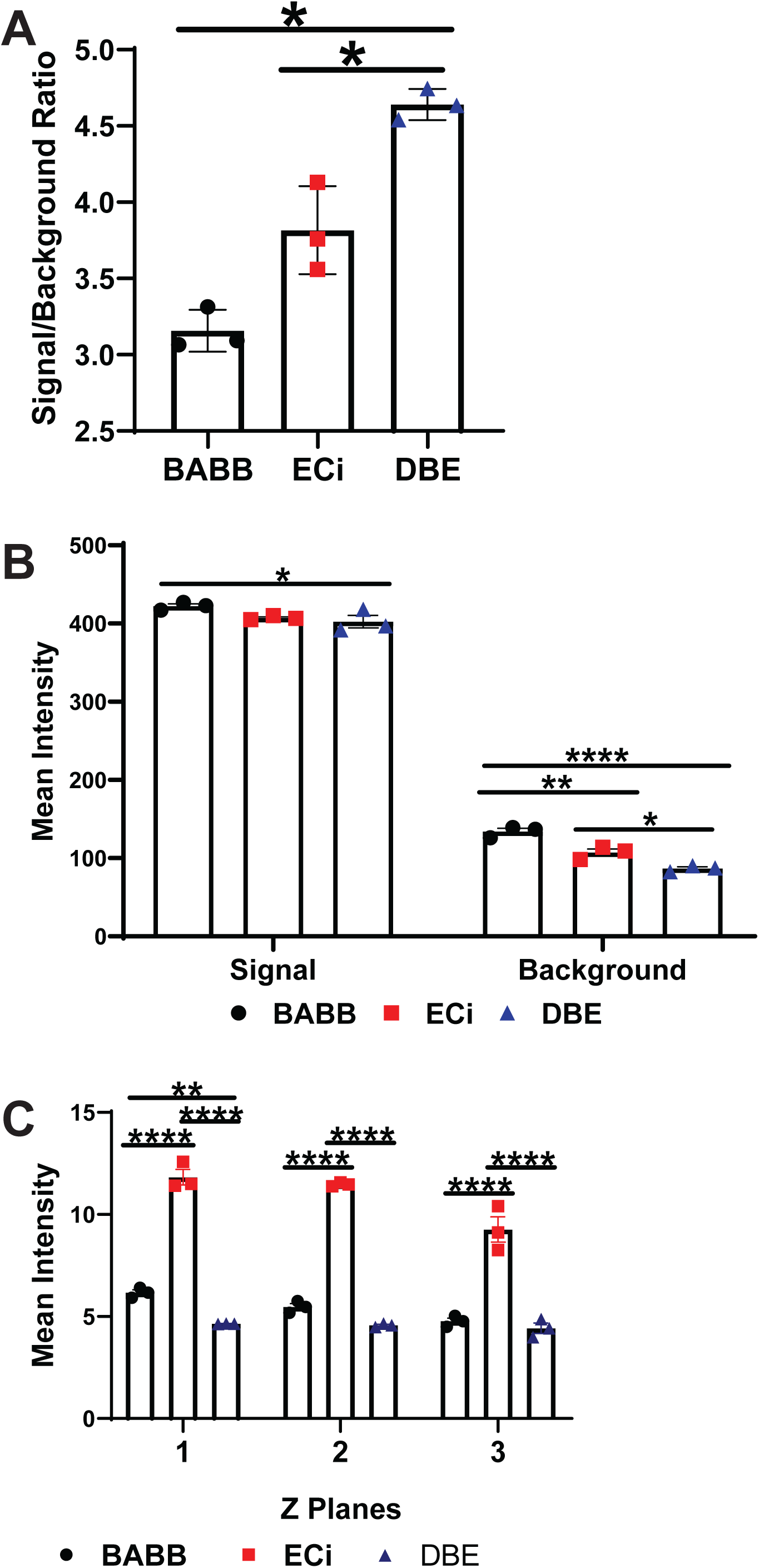
Comparison of different RI imaging media for iDISCO^+^ cleared mouse hindlimb. (**A**) Quantification of signal to background ratio for mouse hindlimbs following perfusion with lectin-649 nm, decalcification with 10% EDTA, iDISCO+ clearing, and RI mathing in either BABB, ECi, or DBE. p ≤0.05, t-test. (**B**) Quantification of mean fluorescence intensity within vessels (i.e. “signal”) and bone autofluorescence (i.e. “background”) for each of the three RI-matching medias. * = p ≤0.05; ** = p ≤0.01; *** = p ≤0.005. t-test. (**C**) Quantification of mean fluorescence intensity in optical sections along the Z-plane (increasing depths from sections 1 to 3). * = p ≤0.05; ** = p ≤0.01; *** = p ≤0.005; **** = p ≤0.001. t-test.

**Figure S4:**
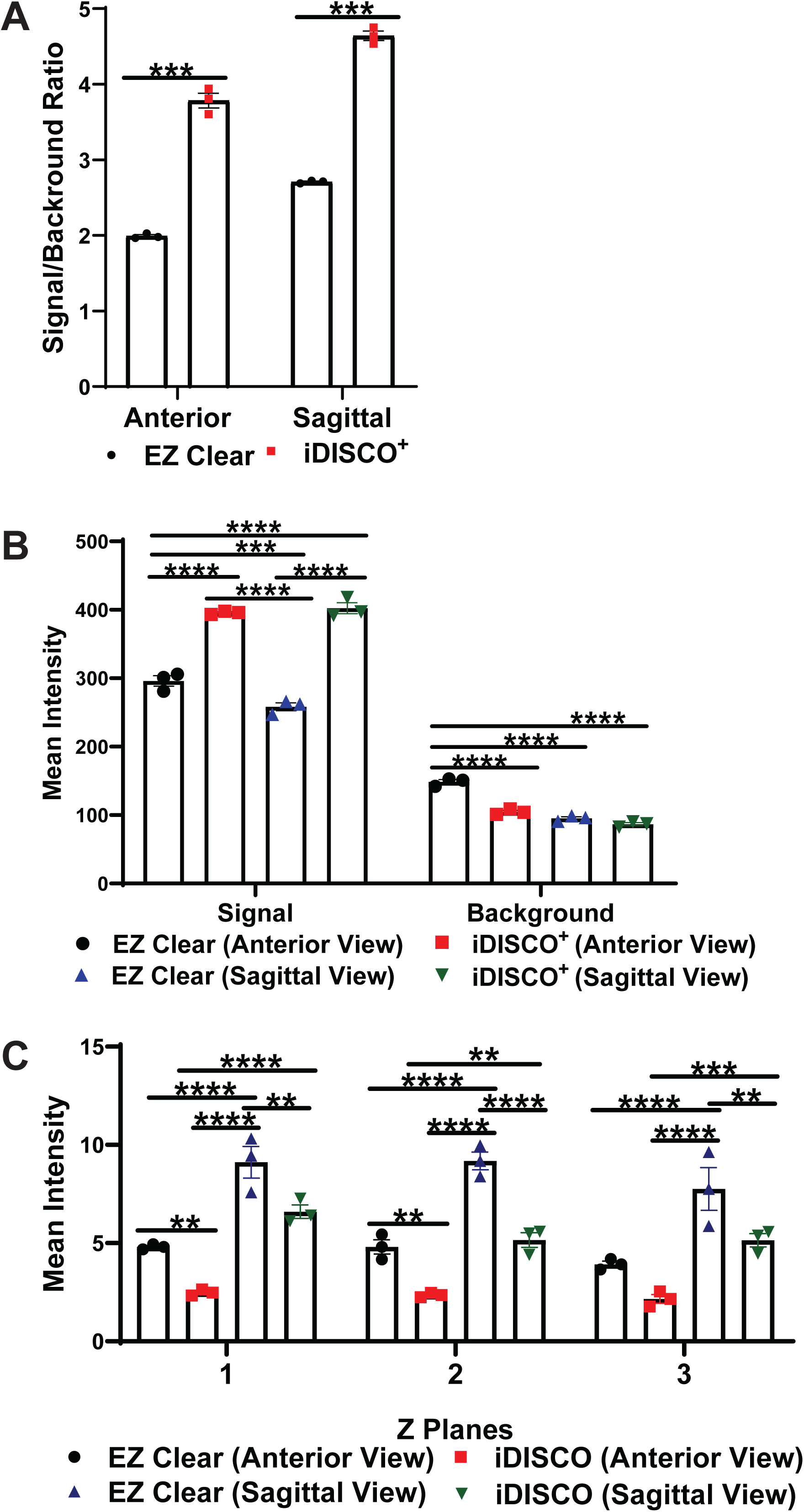
Impact of imaging orientation on image quality for EZ Clear or iDISCO^+^ cleared mouse hindlimbs. (**A**) Comparison of signal to background ratio (SBR) of LSFM images cleared with either EZ Clear or iDISCO^+^ and imaged in different orientations (anterior or sagittal). (**B**) Quantification of lectin-649 fluorescence intensity in vessels (i.e. “signal”) and bone (i.e. “background”). (**C**) Quantification of mean fluorescence intensity in optical sections along the Z-plane (increasing depths from sections 1 to 3). * = p ≤0.05; ** = p ≤0.01; *** = p ≤0.005; **** = p ≤0.001. t-test.

